# Endocytic BDNF secretion through extracellular vesicle-dependent secretory pathways in astrocytes

**DOI:** 10.1101/2024.09.05.611356

**Authors:** Jeongho Han, Hyungju Park

## Abstract

Brain-derived neurotrophic factor (BDNF) plays an essential role in regulating diverse neuronal functions in an activity-dependent manner. Although BDNF is synthesized primarily in neurons, astrocytes can also supply BDNF through various routes, including the recycling of neuron-derived BDNF. Despite accumulating evidence for astrocytic BDNF uptake and resecretion of neuronal BDNF, the detailed mechanisms underlying astrocytic BDNF recycling remain unclear. Here, we report that astrocytic resecretion of endocytosed BDNF is mediated by an extracellular vesicle (EV)-dependent secretory pathway. In cultured primary astrocytes, extracellular BDNF was endocytosed into CD63-positive EVs, and stimulation of astrocytes with ATP could evoke the release of endocytosed BDNF from CD63-positive vesicles. Downregulation of vesicle-associated membrane protein 3 (Vamp3) led to an increase in the colocalization of endosomal BDNF and CD63 but a decrease in extracellular vesicle release, suggesting the necessity of Vamp3-dependent signaling for EV-mediated BDNF secretion. Collectively, our findings demonstrate that astrocytic recycling of neuronal BDNF is dependent on the EV-mediated secretory pathway via Vamp3-associated signaling.

## Introduction

Brain-derived neurotrophic factor (BDNF), a crucial neurotrophin in the brain, is regulated by neuronal activity and plays a crucial role in learning and memory mechanisms^1-3^. BDNF is synthesized primarily in neurons and is secreted in the pro-form of BDNF (proBDNF) or mature form of BDNF (mBDNF)^4^. These secreted BDNF proteins from neurons are essential for long-term synaptic and diverse cognitive functions^5-7^.

Astrocytes also contribute to BDNF-dependent synapse functions. In addition to their own production, astrocytes can also take up extracellular BDNF through the TrkB-T1 receptor^8,9^ and resecrete endocytosed BDNF^10,11^. In our previous study, we reported the uptake of mBDNF via the TrkB receptor and the rerelease of endocytic mBDNF through vesicle-associated membrane protein 3 (Vamp3), a key soluble *N*SF attachment protein receptor (SNARE) protein that is abundant in astrocytes^10^. Because Vamp3 potentially participates in the release of multiple types of vesicles, such as synaptic-like microvesicles (SLMVs), dense-core vesicles (DCVs), and extracellular vesicles (EVs), such as exosomes^12,13^, it is still unclear which vesicular pathway is involved in astrocytic recycling of neuronal BDNF.

Previous studies have revealed that mBDNF and proBDNF, as well as their corresponding receptors, TrkB and p75NTR, are found in EVs^14-19^. These observations suggest that endocytosed BDNF in astrocytes may be secreted through EVs, potentially through Vamp3-dependent signaling. Here, we provide evidence that astrocytes resecrete endocytic BDNF through astrocytic EVs and that Vamp3 is involved in this process. Using quantum dot (QD)-labeled mBDNF (QD-BDNF) as a proxy for extracellular BDNF proteins derived from neurons, we found that endocytosed BDNF is located in CD63-positive vesicles and that ATP-induced endocytic BDNF resecretion from CD63-positive EVs is observed. Additionally, knockdown of Vamp3 resulted in decreased EV release and QD-BDNF secretion from CD63-positive vesicles in astrocytes, indicating that EV-mediated BDNF secretion requires Vamp3-dependent signaling. These results provide insights into a novel mechanism whereby astrocytic BDNF recycling occurs through EVs.

## Results

### Endocytic BDNF in CD63-positive vesicles

In our previous study, we reported that a significant portion of endocytosed BDNF is sorted into Vamp3-positive vesicles in astrocytes, whereas endocytic BDNFs are partially localized in either recycling endosomes or dense-core vesicles^10^. These results suggest that Vamp3-dependent resecretion of endocytic BDNF is mediated by nonclassical recycling or secretory pathways. As Vamp3 has been suggested to be associated with the transport and secretion of EVs^12^, endocytic BDNF in astrocytes may be secreted through EV-dependent secretory pathways.

To explore this possibility, the secretion of endocytic BDNF from cultured primary astrocytes transfected with CD63-EGFP (a marker for EVs) or Rab11-EGFP (a marker for recycling endosomes) was monitored. QD-BDNFs were treated for 30 min or 120 min to detect the inclusion of endocytosed extracellular BDNF and to test whether they were included in the EV or recycling endosome fraction (Fig. 1A). We found that a significant amount of QD-BDNF accumulated in CD63-positive vesicles, and this accumulation increased over time (Fig. 1B). Considerable accumulation of QD-BDNF in Rab11-positive vesicles was also detected, but the level of QD-BDNF in Rab11-positive vesicles was lower than that in CD63-positive vesicles, and QD-BDNF did not accumulate in a time-dependent manner as did QD-BDNF in CD63-positive vesicles (Fig. 1B). These results suggest that CD63-positive vesicles constitute one of the main subcellular locations of endocytosed BDNF in astrocytes.

**Figure 1.**
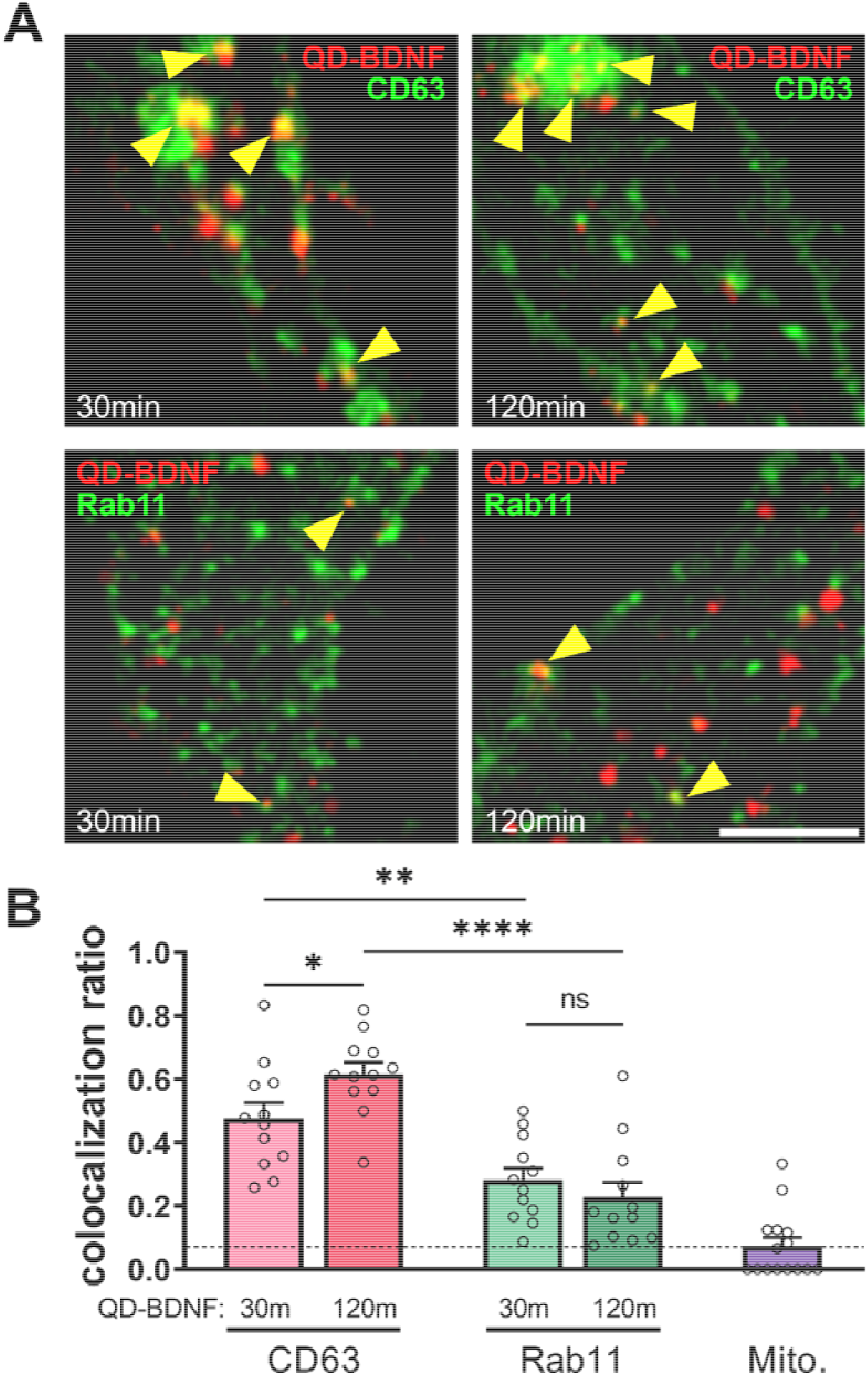
Colocalization of endocytic BDNF with CD63 or Rab11 (A) Representative fluorescence images showing the colocalization of QD-BDNF with CD63 or Rab11 in astrocytes. QD-BDNF was added for 30 or 12 min to cultured astrocytes transfected with CD63-EGFP or Rab11-EGFP. Yellow arrowheads indicate QD-BDNF colocalized with the corresponding vesicular markers. Scale bar = 5 μm. (B) Bar graphs depict average colocalization ratios (number of colocalized QD-BDNF particles among total QD-BDNF particles). Dotted line: average colocalization ratio between QD-BDNF and MitoTracker (Mito.) as a negative control. *\*P* < 0.05. *N* = 12–15 cells (194–2,782 QD particles).

### Astrocytic EVs contain endocytosed BDNF

We next tested whether source cell-derived extracellular BDNF could be found from EVs in astrocytes after endocytosis. After BDNF-EGFP was expressed in differentiated Neuro2a (N2a) cells (Fig. 2A, B), N2a-conditioned medium (NCM) containing BDNF-EGFP secreted from N2a cells was harvested. Then, primary cultured astrocytes were treated with N2a-conditioned media to induce the endocytosis of BDNF-EGFP secreted from N2a cells into astrocytes (Fig. 2A). After being incubated with NCM, astrocytes were treated with ATP (100 µM) to induce the secretion of endocytic BDNF. The astrocyte-conditioned media (ACM) were than collected, and EV fractions were isolated from the harvested ACM (Fig. 2A).

**Figure 2.**
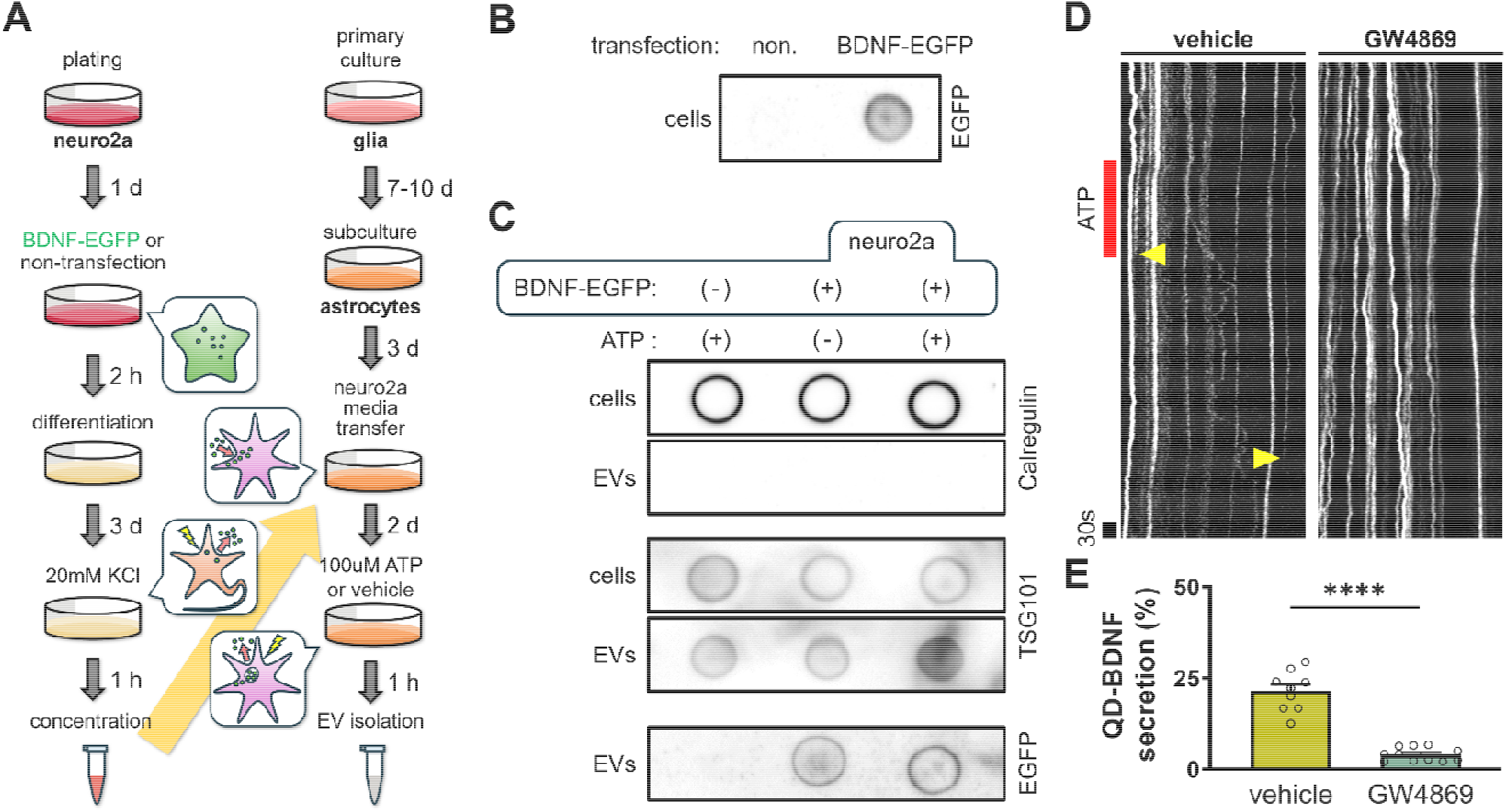
EV-mediated release of endocytosed BDNF from astrocytes (A) Schematic diagram of the experiment for detecting BDNF-EGFP derived from Neuro2A cells in astrocytic EVs. (B) Dot blot showing EGFP expression in cell lysates from nontransfected (non. ; negative control) or BDNF-EGFP-transfected Neuro2a cells. (C) Dot blotting was used to detect protein expression in cell lysates and EVs extracted from the media of astrocytes treated with ATP or vehicle after being supplied with media from nontransfected or BDNF-EGFP-transfected Neuro2a cells. Antibodies targeting general markers of the endoplasmic reticulum (ER; an EV-negative marker, calregulin) or EVs (TSG101) were used. (D) Representative QD-BDNF kymographs from astrocytes were generated via ImageJ/FIJI and obtained after treatment with vehicle or GW4869 (20 μM) for 1 h prior to time-lapse imaging. Red bar: ATP (100 μM) treatment. Arrow heads: disappearance of QD-BDNF fluorescence. (E) Average percentages of ATP-induced QD-BDNF secretion events from vesicles in astrocytes pretreated with either vehicle or GW4869. *\*\*\*\*P* < 0.0001. *N* = 9–11 cells (vehicle = 201; GW4869 = 468 QD particles).

Dot blot analysis of the EV fraction of the harvested ACM revealed the selective presence of TSG101 (EV marker) with an absence of calregulin (ER marker), verifying the EV fraction of ACM (Fig. 2C). In this ACM-derived EV fraction, we successfully detected BDNF-EGFP from ACM harvested from astrocytes treated with NCM derived from N2a cells expressing BDNF-EGFP (Fig. 2C). However, no BDNF-EGFP was detected in EVs collected from astrocytes treated with NCM from nontransfected N2a cells (Fig. 2C). These results suggest that astrocytes release endocytosed BDNF via the EV-mediated secretory pathway.

Further tests support the idea that the resecretion of endocytic BDNF in astrocytes depends on the EV-mediated secretory pathway. We next measured the secretion of endocytic BDNF from astrocytes by monitoring the fluorescence intensity of endocytosed QD-BDNF in cultured astrocytes. In this preparation, the secretion events of intracellular QD-BDNF particles could be detected by the abrupt disappearance of QD-BDNF fluorescence due to the exposure of QD-BDNF to the extracellular QSY21 quencher^10^. GW4869 (20 µM), an inhibitor of EV biogenesis, was used to inhibit EV release from astrocytes. Our results revealed that ATP-induced QD-BDNF secretion was significantly abolished in GW4869-treated astrocytes (Fig. 2D, E).

Together with findings showing significant colocalization of QD-BDNF with CD63-EGFP in astrocytes (Fig. 1), these results indicate that extracellular BDNF originating from other cells, such as neurons, could be sorted to the EV fraction after endocytosis and resecreted via the EV-dependent secretory pathway in astrocytes.

### Vamp3 regulates EV release from astrocytes

Our previous study revealed that endocytosed extracellular BDNF is mainly sorted into Vamp3-positive vesicles and that the resecretion of these endosomal BDNF molecules is dependent on Vamp3-mediated exocytosis^10^. Given that endocytosed BDNF is also found in the EV fraction and is secreted through EV-dependent secretory pathways (Fig. 2), astrocytic release of EVs may require Vamp3 signaling.

To test this possibility, we examined whether the downregulation of astrocytic Vamp3 influences EV release. We assessed the amount of EVs released from cultured astrocytes transfected with scrambled (*siSCR*) or *Vamp3* siRNA (*siVamp3*). Consistent with previous reports showing the facilitation of EV release from glial cells by ATP treatment^20,21^, our data revealed that ATP stimulation (100 µM) increased EV release from astrocytes (Fig. 3A-C).

**Figure 3.**
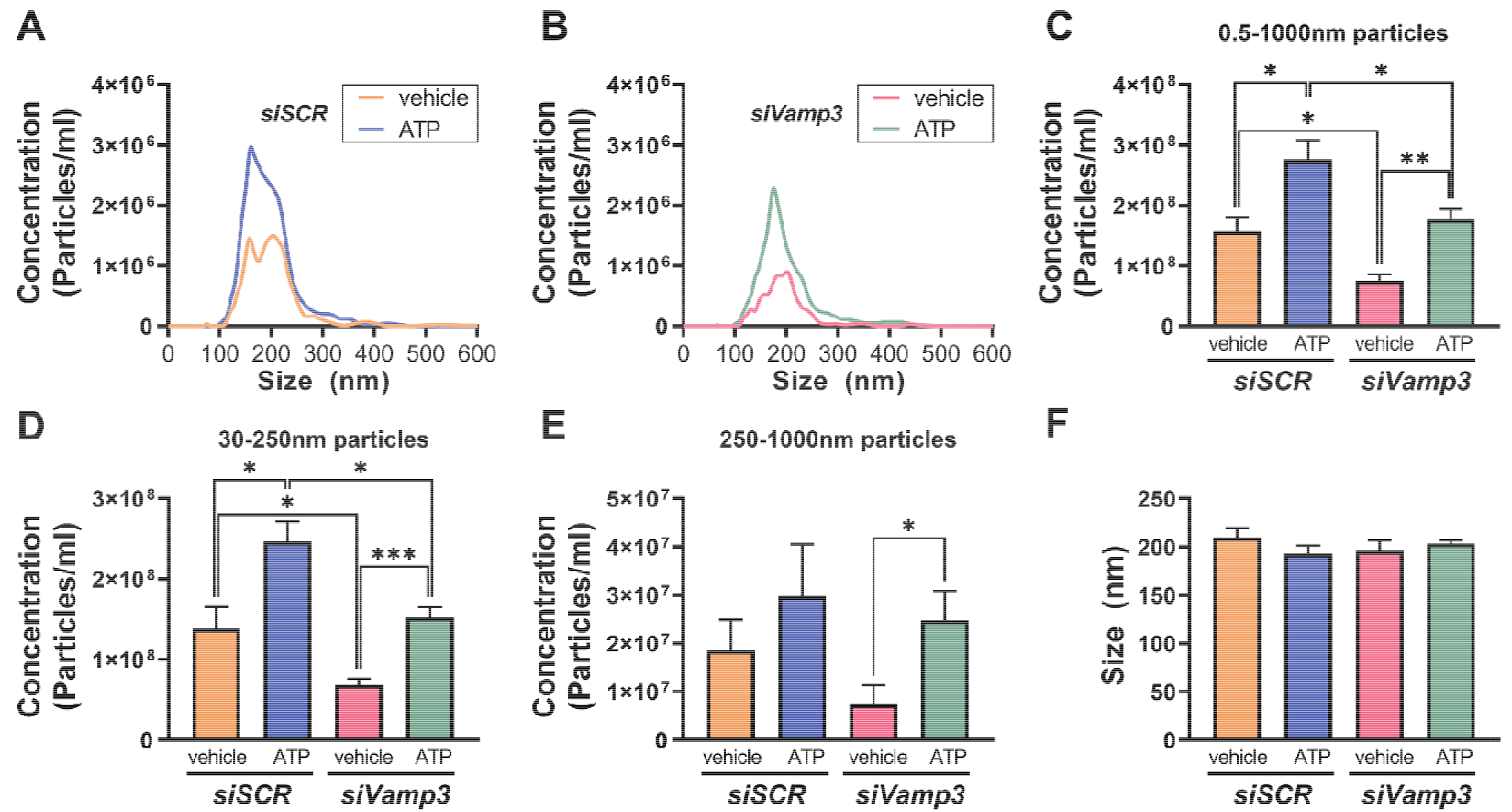
Knockdown of Vamp3 attenuated EV release from astrocytes (A, B) Averaged distribution curves representing the concentrations of particles versus EV sizes isolated from the conditioned media of astrocytes transfected with either *siSCR* or *siVamp3*. Vehicle treatment was used to obtain spontaneously released EVs, while ATP treatment was used to obtain EVs released in response to ATP stimulation. *\*P* < 0.05, *\*\*P* < 0.01, *\*\*\*P* < 0.001. *N* = 5 batches. (C-E) Bar graphs showing the average number of particles categorized by size range. (C): 0.5–1000 nm, (D): 30–250 nm, (E): 250–1000 nm. *\*P* < 0.05, *\*\*P* < 0.01, *\*\*\*P* < 0.001. (F) Bar graphs showing the average particle size from each condition.

When the size of the EV particles was analyzed, ATP stimulation efficiently promoted the release of EVs 30∼250 nm in size (Fig. 3D), whereas the secretion of EVs greater than 250 nm in size was not affected by ATP treatment (Fig. 3E). Whereas spontaneous release of EV particles was also observed (vehicle groups in Fig. 3), the amount of spontaneously released EVs was less than that of ATP-evoked EVs (Fig. 3A-C). The overall average sizes of spontaneous and ATP-evoked EV particles were similar (Fig. 3F).

We found that the number of either spontaneously released or ATP-evoked EV particles was significantly decreased by reduced Vamp3 expression (Fig. 3C). This Vamp3 dependency of spontaneous or ATP-evoked EV release was prominent in EVs 30∼250 nm in size (Fig. 3D), although the reduction in Vamp3 expression resulted in only a slight decrease in the secretion of EV particles larger than 250 nm (Fig. 3E). Our results are in accordance with those of previous reports showing that neuronal EV release is dependent on Vamp3, which can promote the fusion of EV-containing multivesicular bodies (MVBs) to the plasma membrane^22^. Moreover, we found no direct interaction between CD63 and Vamp3 (Fig. S1) and no effect of Vamp3 knockdown (KD) on EV particle size (Fig. 3F), suggesting that the reduction in ATP-evoked EV release caused by Vamp3 depletion is not due to alterations in endosome–EV sorting or trafficking.

### Vamp3 regulates EV-mediated BDNF release from astrocytes

On the basis of the presence of endocytic BDNF in secreted EV fractions (Fig. 2), the requirement of Vamp3 for EVs (Fig. 3) and the release of endocytic BDNF^10^, we hypothesized that Vamp3 regulates the secretion of endocytic BDNF-containing EVs from astrocytes. To address this possibility, we directly tested whether endocytic BDNF secretion from CD63-positive vesicles is altered by reduced Vamp3 expression.

After loading QD-BDNF in CD63-EGFP-expressing astrocytes with *siSCR* or *siVamp3* expression, real-time endocytic QD-BDNF release was monitored by detecting the disappearance of QD-BDNF fluorescence due to the exposure of QD-BDNF to the QSY21 quencher via opened vesicle pores (Fig. 4A)^10^. Consistent with our previous report^10^, ATP stimulation successfully evoked QD-BDNF release from astrocytes. Further analysis revealed that 66.6 ± 3.7% of ATP-induced QD-BDNF secretion was from CD63-positive vesicles (Fig. 4B), indicating that a major population of endocytic BDNF is released through CD63-positive vesicles. The amount of endocytic BDNF released from CD63-positive vesicles was significantly reduced by Vamp3 KD (Fig. 4C). Endocytic BDNF release from CD63-negative vesicles was also downregulated by Vamp3 KD (Fig. 4C), suggesting that a minor population of endocytic BDNF in CD63-negative vesicles also requires Vamp3 for resecretion.

**Figure 4.**
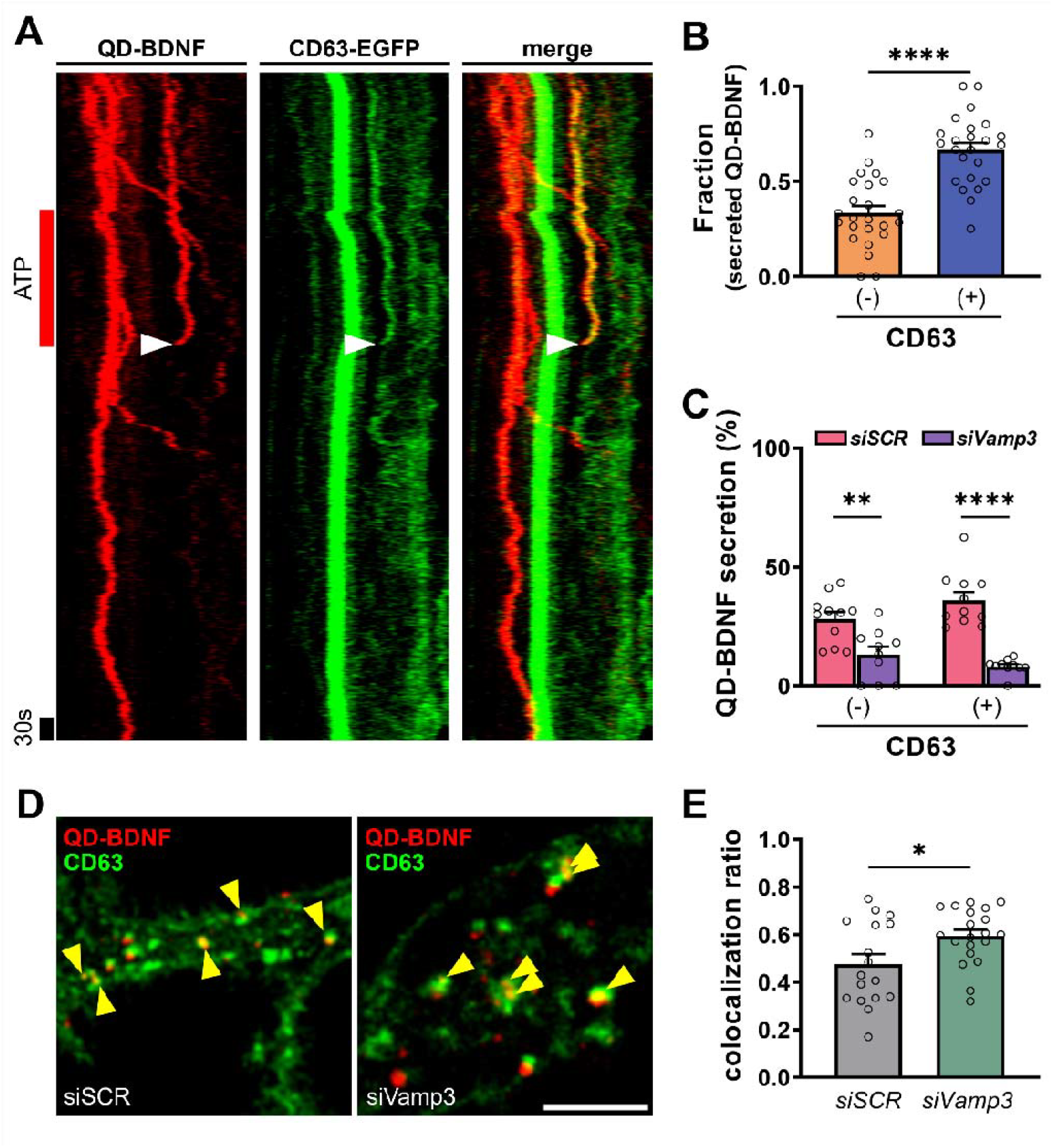
Vamp3-dependent secretion of QD-BDNF from CD63-positive vesicles (A) Representative QD-BDNF kymographs generated from CD63-EGFP-transfected astrocytes. Arrow eads: disappearance of CD63-EGFP-positive QD-BDNF fluorescence. (B) Average fractions of secreted QD-BDNF particles with (+) or without CD63 (-) among total secreted QD-BDNF particles. *N* = 24 cells (232 secreted QD particles). (C) Average percentages of ATP-induced QD-BDNF secretion events from vesicles with (+) or without CD63 (-). *\*\*P* < 0.01, *\*\*\*\*P* < 0.0001. *N* = 10–12 cells (*siSCR* = 508; *siVamp3* = 308 QD particles). (D) Representative fluorescence images showing the colocalization of QD-BDNF with CD63-EGFP in astrocytes treated with *siSCR* or *siVamp3*. Yellow arrowheads indicate QD-BDNF colocalized with CD63-EGFP. Scale bar = 5 μm. (E) Average colocalization ratios (number of colocalized QD-BDNF particles among total QD-BDNF particles). *\*P* < 0.05. *N* = 12 cells (1,737–1,962 QD particles).

To test whether Vamp3 is involved in the transport of endocytic BDNF to the EV fraction, we assessed the effect of Vamp3 KD on the colocalization of QD-BDNF and CD63-EGFP (Fig. 4D). Our results revealed that the colocalization ratio between CD63 and QD-BDNF was significantly elevated by Vamp3 KD (Fig. 4E). Given that Vamp3 KD does not influence overall endocytosis and transport of endocytic BDNF^10^ but reduces EV release (Fig. 3), the Vamp3 KD-induced increase in colocalization between QD-BDNF and CD63 appears to involve the overaccumulation of QD-BDNF-containing EVs in MVBs due to the attenuated docking of EV-containing MVBs to the plasma membrane^22^.

## Discussion

In this study, we demonstrated that astrocytes release endocytosed BDNF via an EV-dependent secretory pathway. Our previous work revealed that Vamp3 is required for ATP-induced release of endocytic BDNF in astrocytes, but the exact secretory pathways responsible for endocytic BDNF release from astrocytes remain elusive.

Our results revealed that up to 60% of endocytosed BDNF could be sorted into CD63-positive EVs after endocytosis (Fig. 1). While both late endosomes and lysosomes are the main endosomal fractions containing CD63, endocytosed BDNF seems not to be directed to these vesicular fractions, as endocytosed QD-BDNF was less targeted to Rab11- or Lamp1-positive vesicles than to Vamp3-positive vesicles^10^. The MVB is another major vesicular fraction containing CD63 and small endosome-derived EVs such as exosomes^23^. Therefore, we hypothesized that the EV fraction is one of the major endocytic/recycling pathways for processing endocytosed BDNF in astrocytes. In support of this idea, we found that endocytic BDNF in astrocytes could be released via the EV-mediated pathway (Fig. 2).

A recent study revealed that Vamp3 plays a role in EV secretion in neurons by facilitating membrane docking and fusion of EV-containing MVBs^22^. Consistent with this finding, our study also revealed a dependency of astrocytic BDNF-containing EV release on Vamp3 (Fig. 4). Vamp3 does not affect the uptake of extracellular BDNF or the overall transport of BDNF-containing endosomes^10^ but induces the overaccumulation of endocytic BDNF in CD63-positive vesicles (Fig. 4D, E) due to the reduced release of BDNF-containing EVs. Therefore, in addition to endosome recycling^24^, our data indicate that astrocytic Vamp3 is strongly implicated in controlling EV secretion as a V-SNARE.

Our study suggested that Vamp3-mediated EV secretion is the major secretory pathway mediating astrocytic rerelease of endocytic BDNF, but other molecular mechanisms involved in astrocytic BDNF recycling cannot be excluded. We found that ATP stimulation, which is known to increase the release of EVs^25^, could elicit endocytic QD-BDNF release not only from CD63-positive EVs but also from CD63-negative vesicles (Fig. 4B). In addition, a reduction in Vamp3 expression also resulted in decreased secretion of endocytic BDNF from both CD63-positive and -negative vesicles (Fig. 4C). On the other hand, our previous report also revealed that Vamp3-positive vesicles constitute the major fraction where endocytic BDNF secretion occurs, whereas a portion of endocytic BDNF is secreted from Vamp3-negative vesicles^10^. Together, these data suggest that multiple endocytic pathways and related mechanisms may be involved in astrocytic recycling of endocytic BDNF. However, further studies are needed to elucidate the unknown mechanisms involved in endocytic BDNF release from astrocytes.

The physiological functions of EV-mediated BDNF release from astrocytes are still unclear. It has been reported that astrocytic rerelease of neuronal proBDNF can contribute to long-term synaptic plasticity and memory formation via activation of neuronal TrkB^11^, but BDNF release via astrocytic EVs is unlikely to be involved in this function. Rather, BDNF, which is directly secreted in an EV-independent manner (Fig. 4B), may diffuse to and activate TrkB receptors in neighboring cells. Given that BDNF-loaded EVs are not limited to local interactions but can travel long distances^26,27^, BDNF-containing EVs from astrocytes may increase BDNF concentrations or trigger synchronized BDNF-dependent signaling pathways in target cell populations. Further investigations of the functional roles of astrocytic EV-mediated BDNF release and its implications in cognitive functions and brain disorders are needed.

## Methods

### Ethical approval

All animal experimental procedures were conducted in accordance with the guidelines and regulations approved by the Institutional Animal Care and Use Committee of the Korea Brain Research Institute (IACUC-2023-00008).

### Primary astrocyte and neuron culture

We used a previously reported AWESAM protocol^28^ with minor modifications to culture astrocytes that exhibited an in vivo-like morphology. Cortical astrocytes were prepared from wild-type C57BL/6 mice at postnatal day 2. The cortices were dissected in dissection medium (10 mM HEPES in HBSS) at 4 °C, followed by incubation in 0.25% trypsin-EDTA at 37 °C for 20 min. After trypsinization, the tissue was washed five times in dissection medium at 4 °C and then triturated with 5 ml of NB+ medium (2% B-27 supplement, 2 mM GlutaMax, 5000 U/ml penicillin, and 5000 μg/ml streptomycin in neurobasal medium). The dissociated cells were filtered through a cell strainer and plated on 0.04% polyethylenimine (PEI)-coated cell culture dishes (3 × 10^6^ cells/150 mm dish) in culture media (10% FBS, 5,000 U/ml penicillin, and 5 mg/ml streptomycin in DMEM). Seven days postplating, the dissociated cells were shaken at 110 rpm for 6 h. Subsequently, the cells were washed three times with 1X PBS, treated with 0.25% trypsin, and plated on 0.04% PEI-coated plates in NB+ medium supplemented with HBEGFP (50 μg/ml). For immunoprecipitation, we utilized a 6-well plate (5 × 10^5^ cells/well), while a 150 mm dish (5 × 10^6^ cells/dish) was used for EV isolation, and 18 mm coverslips in a 12-well plate (1 × 10^5^ cells/well) and glass-bottom dishes (6 × 10^4^ cells/dish) were used for imaging.

Cortical neurons were prepared from embryonic day 16 C57BL/6 mouse embryos. The cortices were dissected in dissection medium (10 mM HEPES in HBSS) at 4 °C, followed by incubation in 0.25% trypsin-EDTA at 37 °C for 20 min. After trypsinization, the tissue was washed five times in dissection medium at 4 °C and then triturated with 5 ml of plating medium (10% FBS, 2 mM glutamine, 25 mM glucose, 5,000 U/ml penicillin, and 5,000 μg/ml streptomycin in MEM). After trituration, the cells were plated in a poly-L-lysine-coated 150 mm dish (1.5 × 10^7^ cells/dish) containing plating medium, and after 2 h, the medium was replaced with maintenance medium (2% B-27 supplement, 2 mM glutamine, 5,000 U/ml penicillin, and 5 mg/ml streptomycin in neurobasal medium).

For Neuro2a differentiation, the cells were first cultured in DMEM supplemented with 10% FBS, 5000 U/ml penicillin, and 5000 μg/ml streptomycin for 1 d. After washing three times with PBS, the cells were then cultured in Opti-MEM supplemented with 0.01% FBS, 5000 U/ml penicillin, and 5000 μg/ml for 3 d.

### Transfection and QD imaging

DNA and siRNA constructs were transfected with Lipofectamine 2000 into Neuro2a or cultured astrocytes at 10–11 DIV according to the manufacturer’s protocol. The knockdown efficiency of Vamp3 was validated in a previous report^10^. QD-BDNF synthesis and QD imaging were performed as previously reported^10^. In brief, 50 nM biotinylated mature BDNF (bt-BDNF) was incubated with 50 nM streptavidin-conjugated quantum dot 655 (st-QD655) at 4 °C overnight at a ratio of 2:1. Subsequently, QD-BDNF was filtered with a 100 kDa Amicon filter and then eluted with 1% BSA in PBS.

For colocalization experiments, astrocytes transfected with CD63-EGFP for 2 days were treated with QD-BDNF for 30 min and then fixed with 4% paraformaldehyde (PFA). For time-lapse imaging, QD-BDNF-treated cells were recorded in an extracellular solution (in mM; 119 NaCl, 2.5 KCl, 20 HEPES, 2 CaCl_2_, 30 glucose, and 2 MgCl_2_, pH 7.4) containing 4 μM QSY21 to quench the extracellular QD signal by using a confocal laser scanning microscope (TCS SP8, Leica) at a 1 Hz rate. To induce the secretion of endocytic QD-BDNF, 100 μM ATP was used for 3 min. QD655 fluorescence was excited via a 561 nm laser and detected with a HyD (hybrid) detector set to a range of 650–695 nm. Before conducting all the experiments with QD-BDNF, quantum dot 655-conjugated bovine serum albumin (QD-BSA) was used as a negative control to confirm the absence of nonspecific QD signals.

### EV isolation

For EV isolation, the procedure was conducted with minor modifications to a published protocol^29^, using two 150 mm dishes of cultured mouse astrocytes. The astrocyte medium was sequentially centrifuged at 4 °C, first at 500 × g for 5 min, then at 2000 × g for 10 min, and finally at 10000 × g for 30 min. At each step, only the supernatant was transferred. After centrifugation, the media was filtered through a 0.22 μm filter, and an equal volume of 16% PEG with 1 M sodium chloride (final PEG concentration: 8%, final sodium chloride concentration: 0.5 M) was added. The mixture was thoroughly mixed by inversion and then incubated overnight at 4 °C. Next, the mixture was centrifuged at 5000 × g for 2 h at 4 °C, and the pellet was suspended in 40 ml of DPBS. The mixture was subsequently ultracentrifuged at 120,000 × g for 90 min at 4 °C, followed by resuspension in 60 μl of PBS or lysis buffer (0.1% Triton X-100 in PBS with proteinase inhibitors). The resuspension was achieved by shaking (250 rpm) at room temperature for 30 min.

### Dot blotting

The nitrocellulose membrane was spotted with 5 μl of protein sample, dried for 20 min, and then respotted with another 5 μl at the same location, followed by an additional drying period for 1 h. The membrane was blocked with 5% skim milk in TBS-T for 1 h and then incubated overnight at 4 °C with anti-EGFP antibody (1:2000). After incubation, the membrane was washed with TBS-T for 30 min, followed by incubation with the secondary antibody at room temperature for 1 h. Subsequently, the membrane was washed with TBS-T for 30 min, and the immunoreactive proteins were detected via an enhanced chemiluminescence (ECL) kit.

### Nanoparticle tracking analysis

Nanoparticle tracking analysis (NTA) was performed via NanoSight LM10 (Malvern Instruments) according to the manufacturer’s user manual. The samples were diluted in PBS to a final volume of 1 mL and then injected into the laser chamber. Particles were detected via a 488 nm laser and a CMOS camera, with each sample recorded for 60 seconds in triplicate, and the resulting data were averaged. Prior to each experiment, PBS was used as a negative control to ensure the absence of particles.

### Coimmunoprecipitation and western blotting

To determine the interaction between CD63 and Vamp3, cultured astrocytes were transfected with CD63-V5 and Vamp3-3xflag via Lipofectamine 2000 at 10–11 DIV according to the manufacturer’s protocol. Forty-eight hours after transfection, the cells were homogenized in immunoprecipitation (IP) lysis buffer (1% Triton X-100 in PBS with protease inhibitor). The total cell lysates were incubated with 2 μg of anti-flag antibody or mouse IgG1 control antibody at 4 °C overnight and then incubated with protein G-agarose for 7 h. The samples were washed with IP lysis buffer and then eluted for 3 min at 95 °C with SDS□PAGE sample buffer. The samples were analyzed via Western blotting with an anti-FLAG antibody (1:5,000) or anti-V5 antibody (1:2,000) and an HRP-conjugated antimouse or anti-rabbit secondary antibody (1:10,000).

### Endogenous neuronal BDNF detection in astrocytes

Mouse cortical neurons were subjected to Lenti-GFAP-TurboID-KDEL (LV-TurboID) or Lenti-GFAP-EGFP (LV-EGFP) transduction with or without biotin at 8 DIV. After 6 days, 20 mM KCl was applied to the neurons for 1 h to induce the secretion of neuronal proteins. The neuron-conditioned media were then concentrated and incubated with astrocytes for 1 h. Subsequently, the astrocytes were fixed with 4% PFA. For immunocytochemistry, the astrocytes were permeabilized with 0.1% Triton X-100 for 10 min and then blocked with 10% normal goat serum and 3% normal donkey serum for 1 h at room temperature. After blocking, the cells were incubated with anti-BDNF (1:50) at 4 °C overnight and then incubated with anti-Alexa 405 secondary antibody (1:100) and anti-streptavidin Alexa 680 (1:100) for 1 h at room temperature.

### Image and statistical analyses

To analyze the release of QD-BDNF particles, regions of interest (ROIs) in astrocytic processes were manually selected and linearized. The linearized time-lapse images were converted into kymographs via the KymographBuilder plugin in ImageJ/FIJI, and exocytosis events were defined by the complete disappearance of QD-BDNF fluorescence. For colocalization analysis of QD-BDNF with CD63-EGFP or Rab11-EGFP, all the observed QD-BDNF particles were converted to binary images and set as ROIs. The colocalization of each QD-BDNF ROI with EGFP signals was analyzed by using methods and formulas detailed in the previous study^10^. To detect endogenous neuronal BDNF in astrocytes, the colocalization of BDNF and streptavidin signals was analyzed via the colocalization threshold plugin in ImageJ/FIJI, which automatically measures the Manders overlap coefficient^30^. Data are presented as a proportion of the total. Statistical analyses were performed via Prism 8.0 software (GraphPad). Statistically significant differences between two groups were determined via an unpaired Student’s t test, whereas comparisons among three or more groups were made via one-way ANOVA followed by Dunnett’s multiple comparisons test. The Kolmogorov□Smirnov test was used to assess the statistical significance of cumulative distribution percentages between two groups. All the data are presented as the mean ± standard error of the means (SEMs) from three independent batches.

**Table 1.**
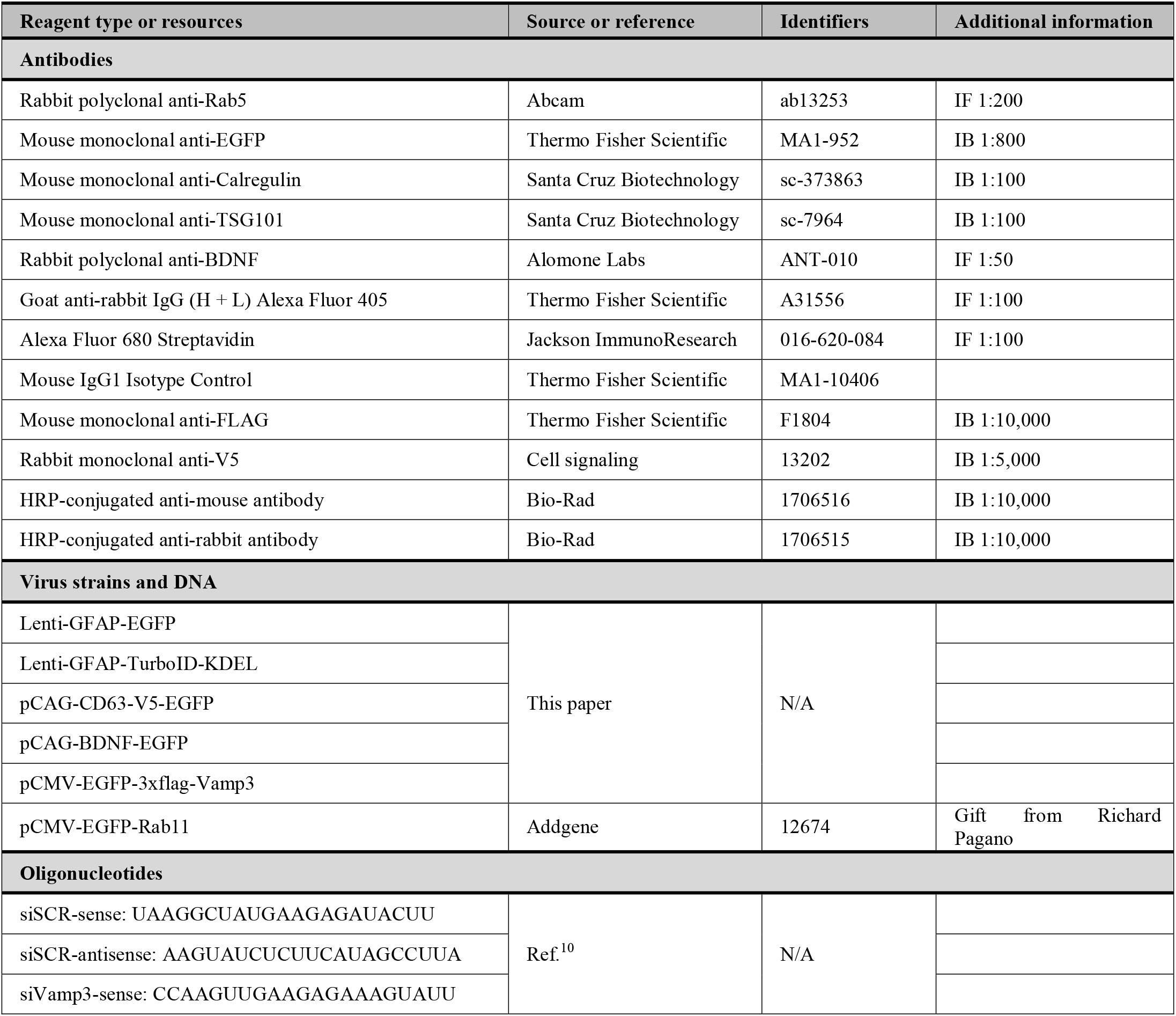
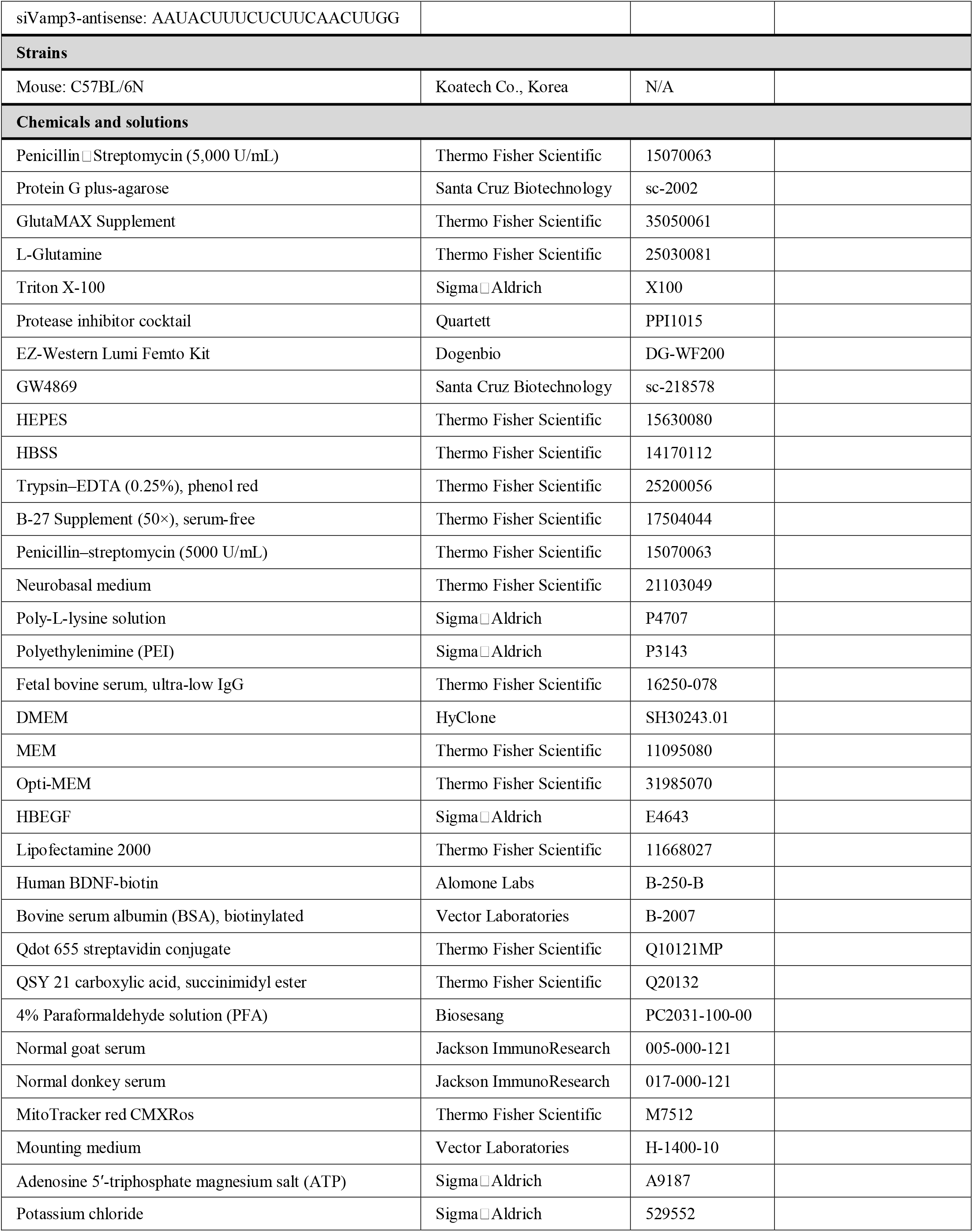
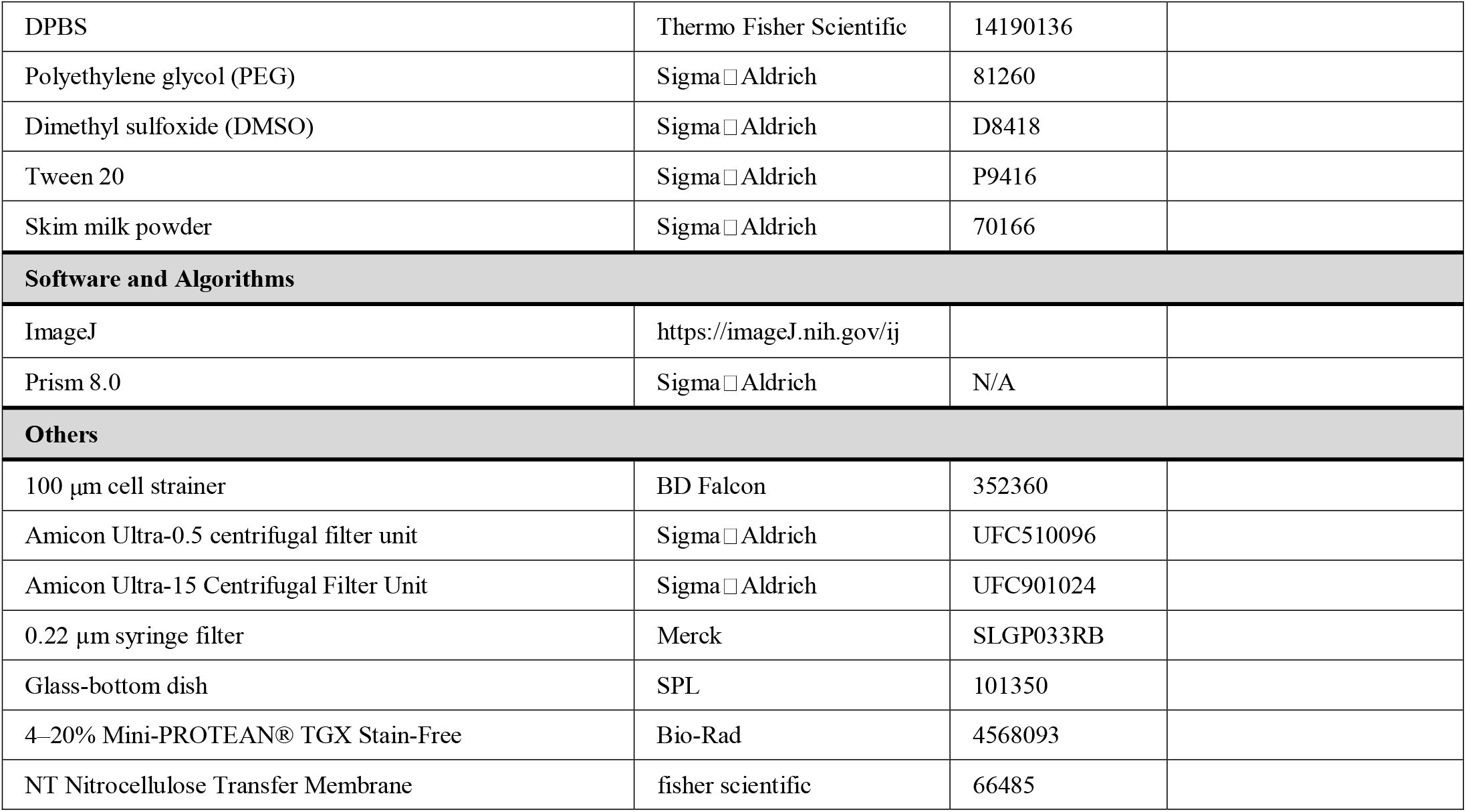
Key resource table. *HRP*, horseradish peroxidase; *GW4869, N,N*′-Bis[4-(4,5-dihydro-1H-imidazol-2-yl)phenyl]-3,3′-p-phenylene-bis-acrylamide; *HEPES*, hydroxyethyl piperazine ethane sulfonic acid; *HBSS*, Hank’s balanced salt solution; *DMEM*, Dulbecco’s modified Eagle’s medium; *MEM*, minimum essential medium; *HBEGF*, heparin-binding EGF-like growth factor; *DPBS*, Dulbecco’s phosphate-buffered saline.

## Acknowledgments

This research was supported by the KBRI basic research program through the Korea Brain Research Institute funded by the Ministry of Science and ICT (24-BR-01-02).

## Author contributions statement

J.H. and H.P. conceived the experiments. J.H. conducted the experiments, analyzed the data, and performed the statistical analysis and figure generation. J.H. and H.P. wrote the manuscript. All the authors reviewed the manuscript.

