## Supplementary Figure 1 for "Endocytic BDNF secretion through extracellular vesicle-dependent secretory pathways in astrocytes"

**Astrocytic BDNF recycling through extracellular vesicle-dependent secretory pathways**

**Supplementary Figure**

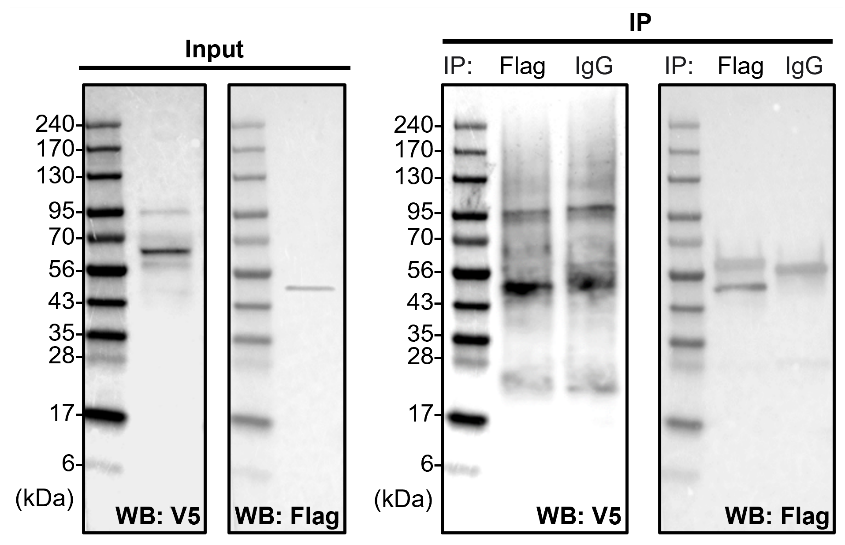

**Figure S1.** Lack of interaction between CD63 and Vamp3 in astrocytes

To investigate the association of CD63-V5 with Vamp3-3xflag, astrocytes were co-transfected with CD63-V5 and Vamp3-3xflag. After 48 hours, immunoprecipitation was conducted using anti-flag or IgG control antibody on the cell lysates. Western blot analysis was then performed using anti-V5 and anti-flag antibodies.
